# Functional correlation of H3K9me2 and nuclear compartment formation

**DOI:** 10.1101/2020.08.28.271221

**Authors:** Kei Fukuda, Chikako Shimura, Hisashi Miura, Akie Tanigawa, Takehiro Suzuki, Naoshi Dohmae, Ichiro Hiratani, Yoichi Shinkai

## Abstract

**Background:** Histone H3 lysine 9 dimethylation (H3K9me2) is a highly conserved silencing epigenetic mark. Chromatin marked with H3K9me2 forms large domains in mammalian cells and correlates well with lamina-associated domains and the B compartment. However, the role of H3K9me2 in 3-dimensional (3D) genome organization remains unclear.

**Results:** We investigated the genome-wide H3K9me2 distribution, the transcriptome and 3D genome organization in mouse embryonic stem cells (mESCs) upon the inhibition or depletion of H3K9 methyltransferases (MTases) G9a/GLP, SETDB1, and SUV39H1/2. We found that H3K9me2 is regulated by these five MTases; however, H3K9me2 and transcription in the A and B compartments were largely regulated by different sets of the MTases: H3K9me2 in the A compartments were mainly regulated by G9a/GLP and SETDB1, while H3K9me2 in the B compartments were regulated by all five H3K9 MTases. Furthermore, decreased H3K9me2 correlated with the changes to the more active compartmental state that accompanied transcriptional activation.

**Conclusion:** Our data showed that H3K9me2 domain formation is functionally linked to 3D genome organization.

## Background

Post-translational modifications of histone proteins regulate chromatin compaction and mediate epigenetic transcriptional regulation. The methylation of histone tails is one of the fundamental events of epigenetic signaling. In organisms ranging from the fission yeast *Schizosaccharomyces pombe* to humans, repeat-rich constitutive heterochromatin is marked by H3K9me2 or H3K9me3 [1–3]. These modifications are catalyzed by a family of SET domain-containing lysine methyltransferases, of which five are present in mammals. SETDB1 and the related enzymes, SUV39H1 and SUV39H2, contribute to the formation of H3K9me3 [1, 4], while GLP and G9a (also called EHMT1 and EHMT2, respectively) regulate H3K9me1 and H3K9me2 formation, respectively [5–7]. In mESCs, H3K9me3 is enriched in retroelements and pericentromeric satellite repeats that are mediated by SETDB1 and SUV39H1/2, respectively [8–10]. SUV39Hs are also involved in H3K9me3 modification in intact retroelements and LINE1 [11]. Unlike H3K9me3, H3K9me2 forms mega base-scale domains that comprise approximately half of the genomes of pluripotent and differentiated cells [9, 12, 13]. Endogenous G9a and GLP mostly exist as a G9a/GLP heterodimer, and the G9a/GLP heteromeric complex is the functional form for global H3K9 methylation *in vivo* [5, 7].

Although H3K9me2 is primarily catalyzed by G9a/GLP in mESCs, not all of H3K9me2 is diminished in either *G9a* or *Glp* knockout (KO) or *G9a/Glp* double knockout (DKO) mESCs [6, 7], which suggests the involvement of other histone MTases in H3K9me2. However, the function and mechanism of G9a/GLP-independent H3K9me2 are currently unknown. H3K9me2 is enriched in pericentromeric satellite repeats [6, 14] and genomic regions with H3K9me3 [15], and SETDB1 and SUV39H1/2 may be involved in G9a/GLP-independent H3K9me2.

Recent Hi-C technology has revealed that chromosomes are hierarchically folded at different levels in the nucleus, such as chromatin loops, topologically associating domains (TADs), and genomic compartments [16–18]. At the mega-base scale, genomic regions are spatially segregated into two subnuclear compartments that predominantly consist of either euchromatin (the A compartment) or heterochromatin (the B compartment). The B compartment is correlated with lamina-associated domains (LADs) [19, 20], which interact with the nuclear lamina (NL) and highly overlap with the H3K9me2 domains [13]. G9a-mediated H3K9me2 is required for LAD-nuclear lamina interaction [21].

Based on the links among H3K9me2, LADs, and B compartments, both G9a/GLP-dependent and independent H3K9me2 could be involved in the A/B compartment formation.

In this study, we analyzed H3K9me2 genome-wide profiles and 3D genome organization in *Setdb1* KO and/or *Suv39h1/2* DKO cells treated with a G9a/GLP specific inhibitor, UNC0642 [22], to identify the role of each H3K9 MTase in H3K9me2 domain formation and whether H3K9me2 is involved in nuclear compartment formation.

## Results

### Compartment-dependent regulation of H3K9me2 in mESCs

It has been reported that H3K9me2 is enriched in LADs, which significantly overlap with the B compartments in mESCs. Although H3K9me2 chromatin immunoprecipitation sequencing (ChIP-seq) in mESCs showed higher H3K9me2 enrichment in the B compartments (negative score regions) than in the A compartments (positive score regions), as expected, H3K9me2 was also present in the A compartments, with a relatively low compartment score (Fig. 1A). As SETDB1 and SUV39H1/2 have the potential to mediate H3K9me2, H3K9me2 ChIP-seq was performed in previously established *Setdb1* conditional KO mESCs[8] or *Suv39h1/2* DKO mESCs[10] (Additional file 1: Fig. S1A) to investigate the role of SETDB1 and SUV39H1/2 in H3K9me2 formation. Although the global H3K9me2 profile did not change largely in *Setdb1* KO or *Suv39h1/2* DKO mESCs (Additional file 1: Fig. S1B), H3K9me2 decreased in the A and B compartments in *Setdb1* KO mESCs and *Suv39h1/2* DKO mESCs, respectively (Fig. 1B).

**Fig. 1.**
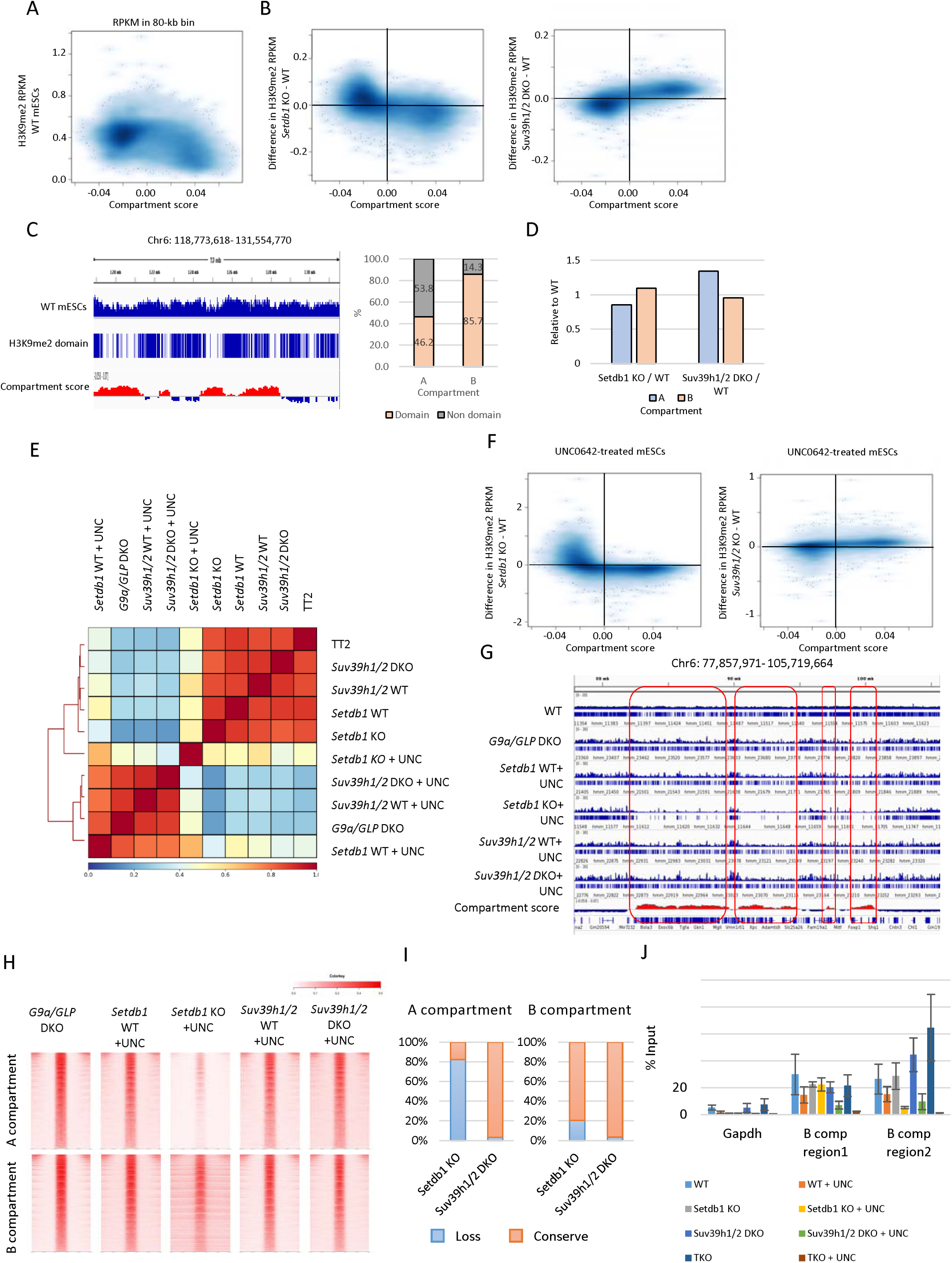
Characterization of Setdb1 and Suv39h1/2-dependent H3K9me2 region in mESCs A. Comparison between compartment score and H3K9me2 enrichment in 80-kb bin. RPKM of H3K9me2 from WT mESCs is negatively correlated with compartment score. Darker blue represents higher dot density. B. Comparison between compartment score in WT mESCs and changes in H3K9me2 in *Setdb1* KO (left) or *Suv39h1/2* DKO (right) mESCs in 80-kb bins. Decreased H3K9me2 is observed in the A and B compartments in *Setdb1* KO and *Suv39h1/2* DKO mESCs, respectively. C. A representative view of H3K9me2 domains in WT mESCs identified by *Hiddendomains* (left). Top panel, H3K9me2 ChIP-seq data from WT mESCs; middle panel, H3K9me2 domains; bottom panel, compartment score in 80-kb bins. Right, fractions of H3K9me2 domains in the A or B compartments. Although H3K9me2 domains are more enriched in the B compartments than in the A compartments, H3K9me2 occupies about half of the A compartments. D. Length of H3K9me2 domains in each KO mESCs to WT mESCs. The length of H3K9me2 domains in the A compartments is slightly decreased in *Setdb1* KO mESCs, while that is increased in *Suv39h1/2* DKO mESCs. E. A correlation matrix of H3K9me2 ChIP-seq data in mESCs. UNC0642-treated mESCs form a cluster distinct from untreated mESCs. Among UNC0642-treated mESCs, *Setdb1* KO mESCs show a different H3K9me2 profile. F. A comparison between compartment score in WT mESCs and changes in H3K9me2 in *Setdb1* KO (left) or *Suv39h1/2* DKO (right) mESCs treated with UNC0642. Consistent with UNC0642-untreated mESCs, H3K9me2 is decreased in the A and B compartments in *Setdb1* KO and *Suv39h1/2* DKO mESCs treated with UNC0642, respectively. G. A representative view of H3K9me2 ChIP-seq from *G9a/GLP* DKO and UNC0642-treated mESCs. *G9a/GLP* DKO mESCs show an H3K9me2 profile similar to UNC0642-treated mESCs. A drastic loss of H3K9me2 domains in UNC0642-treated *Setdb1* KO mESCs in the A compartments (circled in red). H. Heatmaps of H3K9me2 enrichment around G9a/GLP-independent H3K9me2 regions. The A compartment-specific H3K9me2 loss is observed in UNC0642-treated *Setdb1* KO mESCs. I. Fractions of conserved or lost G9a/GLP-independent H3K9me2 in *Setdb1* KO or *Suv39h1/2* DKO mESCs treated with UNC0642. More than 80% of G9a/GLP-independent H3K9me2 domains in the A compartments are lost in UNC0642-treated *Setdb1* KO mESCs, while about 80% of that in B compartment are conserved. In UNC0642-treated *Suv39h1/2* DKO mESCs, almost all G9a/GLP-independent H3K9me2 domains are maintained. J. H3K9me2 ChIP-qPCR analysis in regions shown in Fig. S1N. ChIP was performed five days after 4-OHT treatment in *Setdb1* KO mESCs and 7 days after 4-OHT treatment in *Setdb1*/Suv39h1/2 TKO mESCs. H3K9me2 in the region 1 and 2 is completely lost in UNC0642-treated TKO mESCs.

To obtain a more detailed view of the H3K9me2 profile, we identified H3K9me2 domains in mESCs using *Hiddendomains*, which is a program that uses a hidden Markov model to identify broad domains [23]. The H3K9me2 domains occupied 46.2% and 85.7% of the A and B compartments, respectively (Fig. 1C). The total length of H3K9me2 domains in the A compartments slightly decreased in *Setdb1* KO mESCs, which indicates that SETDB1 may have a role in H3K9me2 domain formation in the A compartments (Fig. 1D). It is difficult to further dissect SETDB1- and SUV39H1/2-dependent H3K9me2 in each KO mESC owing to the dominant impact of G9a/GLP for H3K9me2 formation. A G9a/GLP catalytic inhibitor, such as UNC0642, is useful for analyzing G9/GLP-independent H3K9me2 [22]. Treatment with 0.5-2 μM UNC0642 for three days decreased H3K9me2 to a level comparable with that of *G9a/GLP* DKO mESCs (Additional file 1: Fig. S1C).

Furthermore, H3K9me2 in *G9a/GLP* DKO mESCs and UNC0642-treated mESCs were enriched in pericentromeric satellite repeats, characterized as DAPI-dense regions (Additional file 1: Fig. S1D, E), which is consistent with a previous finding [14]. Compared to WT mESCs, *G9a/GLP* DKO mESCs and UNC0642-treated mESCs formed relatively narrow H3K9me2 domains, which highly overlapped between *G9a/GLP* DKO mESCs and UNC0642-treated mESCs (Additional file 1: Fig. S1F, G). Therefore, UNC0642 treatment can mimic the H3K9me2 profile associated with the *G9a/Glp* DKO phenotype. To investigate the roles of SETDB1 and SUV39H1/2 in G9a/GLP-independent H3K9me2, we performed H3K9me2 ChIP-seq from *Setdb1* KO or *Suv39h1/2* DKO mESCs treated with 2 μM UNC0642. Hierarchical clustering analysis of the H3K9me2 ChIP-seq data showed that the H3K9me2 profile of UNC0642-treated *Setdb1* KO mESCs was largely distinct from that of the *G9a/GLP* DKO mESCs (Fig. 1E). Consistent with UNC0642-untreated mESCs, reductions of H3K9me2 in *Setdb1* KO and *Suv39h1/2* DKO mESCs treated with UNC0642 were mainly observed in the A and B compartments, respectively (Fig. 1F), and a significant reduction of the H3K9me2 domains in the A compartments was observed in UNC0642-treated *Setdb1* KO mESCs (Fig. 1G, H, Additional file 1: S1H). Approximately 80% of the G9a/GLP-independent H3K9me2 domains in the A compartments, and approximately 20% of those in the B compartments were lost in the UNC0642-treated *Setdb1* KO mESCs, while almost all the G9a/GLP-independent H3K9me2 domains were maintained in UNC0642-treated Suv39h1/2 DKO mESCs (Fig. 1I). Although SUV39H1/2 depletion did not have the same impact as SETDB1 depletion from the view of H3K9me2 ChIP-seq data, H3K9me2 immunofluorescent analysis showed a noticeable loss of H3K9me2 signals in DAPI-dense regions in UNC0642-treated *Suv39h1/2* DKO mESCs (Additional file 1: Fig. S1I). This finding suggests a crucial role of SUV39H1/2 in H3K9me2 in pericentromeric satellite repeats.

To investigate whether H3K9me2 in the B compartment was mediated by SETDB1 and SUV39H1/2 redundantly, we established *Setdb1* and *Suv39h1/2* triple KO (TKO) mESCs using the CRISPR-Cas9 system (Additional file 1: Fig. S1J). The complete removal of *Suv39h1* exon 3, which contains SET domain, and the removal of SUV39H2 H398, which is an essential for the methyltransferase activity, were validated by RNA-seq in TKO mESCs[1] (Additional file 1: Fig. S1J). The depletion of SUV39H1/H2 in the TKO mESCs was further validated by western blotting (Additional file 1: Fig. S1K). We used 4-OHT inducible conditional *Setdb1* KO mESCs as parental cells for the TKO mES line. SETDB1 and H3K9me3 were undetectable seven days after 4-OHT treatment (Additional file 1: Fig. S1L). Furthermore, UNC0642 treatment to 4-OHT-treated TKO mESCs decreased H3K9me2 to an undetectable level, as analyzed using western blot (Additional file 1: Fig. S1M). We confirmed the loss of H3K9me2 in the two regions where H3K9me2 was retained in both *Setdb1* KO and *Suv39h1/2* DKO mESCs treated with UNC0642 (Fig. 1J and Additional file 1: Fig. S1N). In conclusion, G9a/GLP-independent H3K9me2 in the A and B compartments in mESCs was mostly mediated by SETDB1 and by both SETDB1 and SUV39H1/2, respectively.

### Compartment-dependent regulation of H3K9me2 in immortalized mouse embryonic fibroblasts (iMEFs)

G9a/GLP-independent H3K9me2 in the A and B compartments was mediated by SETDB1 and both SETDB1 and SUV39H1/2 in mESCs, respectively. To investigate whether this trend is specific to pluripotent stem cells, we analyzed the H3K9me2 profiles in iMEFs. Similar to mESCs, H3K9me2 was more enriched in the B compartments than in the A compartments in iMEFs (Additional file: Fig. S2A). Compared to mESCs, the larger size of the H3K9me2 domains was retained by UNC0642 treatment: the average size of the H3K9me2 domains was 54.5 kb in iMEFs and 14.2 kb in mESCs (Fig. 2A, B). Interestingly, a marked reduction of H3K9me2 in the A compartments was observed in UNC0642-treated iMEFs but not in the B compartments. H3K9me2 decreased upon UNC0642 treatment in approximately 90.0% of the A compartments (36.5/4.1 + 36.5), but only decreased in approximately 35.7% of the B compartments (21.2/21.2 + 36.5) (Fig. 2A, C). G9a/GLP-independent H3K9me2 domains in iMEFs occupied 80.8% of the B compartments and 31.3% of A the compartments (Fig. 2D). Therefore, it can be said that H3K9me2 in the B compartments was more resistant to UNC0642 treatment than that in the A compartments in iMEFs. Despite the crucial role of G9a/GLP in the A compartments, H3K9me2 was not completely diminished by UNC0642 treatment in iMEFs.

**Fig. 2.**
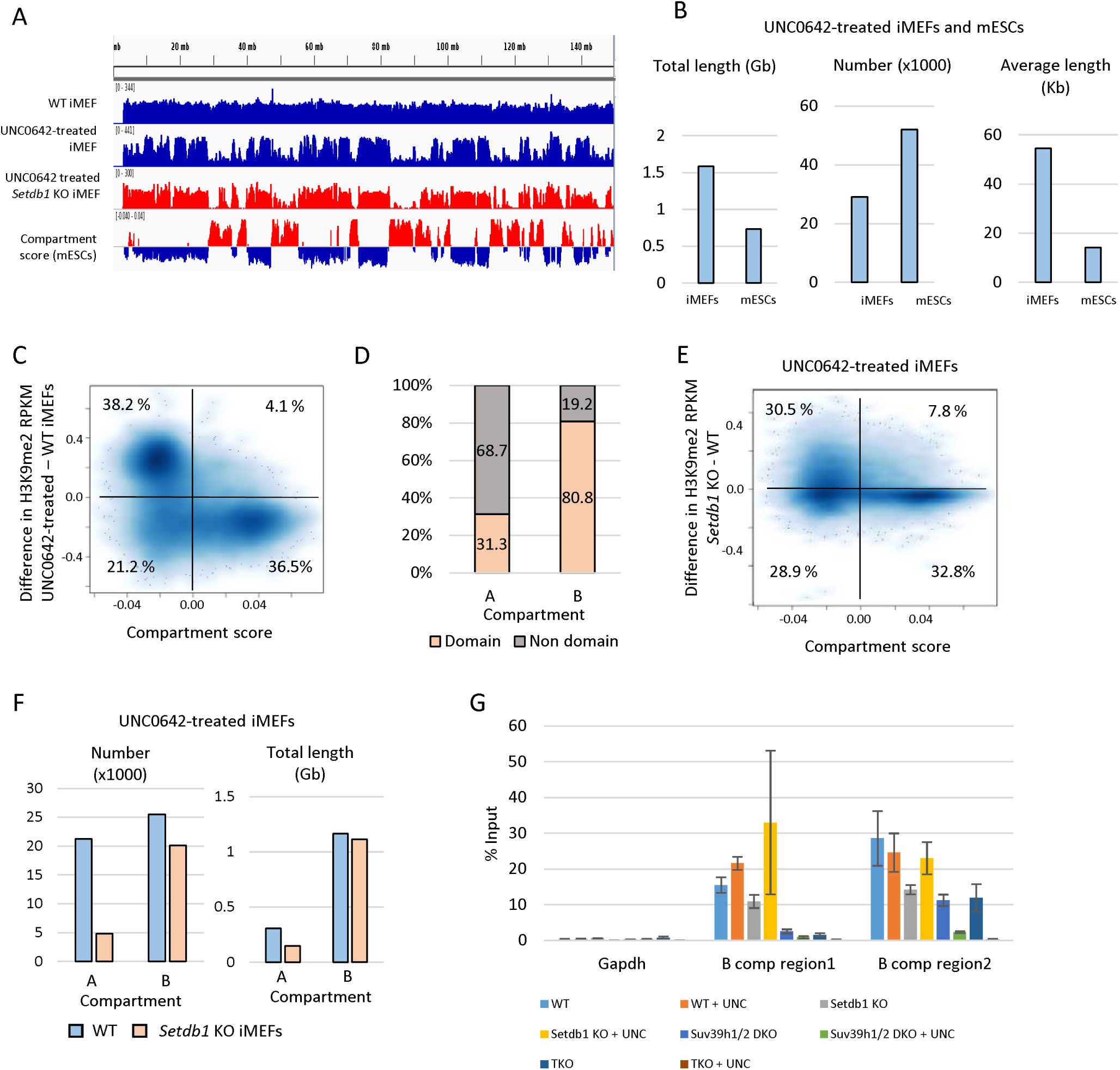
Characterization of Setdb1- and Suv39h1/2-dependent H3K9me2 regions in iMEFs A. A representative view of H3K9me2 ChIP-seq data along chromosome 6 in iMEFs. Although H3K9me2 in A compartment is decreased by UNC0642 treatment, low H3K9me2 signal is retained. Those retained H3K9me2 is further diminished in UNC0642-treated *Setdb1* KO iMEFs. B. Difference in G9a/GLP-independent H3K9me2 between mESCs and iMEFs. H3K9me2 domain size is larger in iMEFs than in mESCs. C. A comparison between compartment score and H3K9me2 changes by UNC0642 treatment in iMEFs. Compartment score in mESCs was used for the analysis. Overall H3K9me2 reduction in the A compartments is induced by the UNC0642 treatment. Each plot represents data from each 80-kb bin. Darker blue represents higher dot density. D. Fractions of H3K9me2 domains in the A and B compartments in UNC0642-treated iMEFs. H3K9me2 domains in the B compartments is largely retained after UNC0642 treatment. E. A comparison between compartment score and H3K9me2 changes in UNC0642-treated *Setdb1* KO iMEFs. Further reduction of H3K9me2 in the A compartments is observed in UNC0642-treated *Setdb1* KO iMEFs. F. The number and the length of H3K9me2 domains in WT and *Setdb1* KO iMEFs treated with UNC0642. Both the number and the length of H3K9me2 domains in the A compartments are reduced in UNC0642-treated *Setdb1* KO iMEFs. G. H3K9me2 ChIP-qPCR analysis in regions showing in Fig. S1N. H3K9me2 in the region 1 and 2 is completely lost in UNC0642-treated TKO iMEFs (N = 3).

We also performed H3K9me2 ChIP-seq analysis on UNC0642-treated *Setdb1* KO iMEFs, which is previously established [24]. Further reduction of H3K9me2 in the A compartments was observed in UNC0642-treated *Setdb1* KO iMEFs (Additional file 1: Fig. S2B). It was observed that 80.8% of the A compartments (32.8/7.8 + 32.8) and 48.7% of the B compartments (28.9/30.5 + 28.9) showed lower H3K9me2 levels in the UNC0642-treated *Setdb1* KO iMEFs than in the UNC0642-treated WT iMEFs (Fig. 2E). The number and total length of the H3K9me2 domains also decreased in the A compartments in the UNC0642-treated *Setdb1* KO iMEFs, while those in the B compartments largely remained unchanged (Fig. 2F). Therefore, SETDB1 is essential for G9a/GLP-independent H3K9me2 in the A compartments in iMEFs, as observed in mESCs.

To investigate whether SETDB1 and SUV39H1/2 mediated all G9a/GLP-independent H3K9me2 in iMEFs, we established *Setdb1* and *Suv39h1/2* TKO iMEFs using the CRISPR-Cas9 system (Additional file 1: Fig. S2C). The established TKO iMEFs showed an undetectable level of H3K9me3, as observed from the results of western blot analysis (Additional file 1: Fig. S2D). H3K9me2 still remained, albeit at a low level, but again became undetectable after UNC0642 treatment in *Setdb1* and *Suv39h1/2* TKO iMEFs (Additional file 1: Fig. S2E). Almost complete loss of H3K9me2 in UNC0642-treated TKO iMEFs was also validated using mass spectrometry analysis (Additional file 1: Fig. S2F). ChIP-qPCR analysis of the selected regions in the B compartments, shown in Additional file 1: Fig. S1N, also showed a loss of H3K9me2 in UNC0642-treated TKO iMEFs (Fig. 2G). Thus, we concluded that H3K9me2 in B compartments is redundantly regulated by G9a/GLP, SETDB1, and SUV39H1/2, both in mESCs and iMEFs.

### G9a/GLP-independent H3K9me2 is correlated with efficient H3K9me2 recovery

Heterochromatin can spread along chromatin from the nucleation sites, such as recruiters of the writer molecule binding regions [25, 26]. As G9a and Glp bind to nucleosomes that contain H3K9me2 via ankyrin repeats of G9a and Glp [27] and methylate the adjacent nucleosome [28], we hypothesized that H3K9me2 spreads from the G9a/GLP-independent H3K9me2 sites during H3K9me2 domain formation. To investigate this, we analyzed H3K9me2 recovery in mESCs after UNC0642 treatment (Fig. 3A). As H3K9me2 almost completely recovered three days after UNC0642 removal (Additional file 1: Fig. S3A), we performed H3K9me2 ChIP-seq at each time point after UNC0642 removal (0h, 24h, 32h, 40h, 48h, 56h, 64h, and 72h). From the H3K9me2 ChIP-seq time course data, we obtained H3K9me2 dynamics during the recovery period (Fig. 3B). Principal component analysis of H3K9me2 Reads Per Kilobase Million (RPKM) in the 80-kb genomic window showed gradual H3K9me2 recovery after UNC0642 removal (Fig. 3C). To investigate the role of G9a/GLP-independent H3K9me2 in H3K9me2 recovery, we classified genomic regions as “early,” “middle,” or “late,” based on the timing of H3K9me2 recovery after UNC0642 removal and compared H3K9me2 levels among the different genomic region classes in the UNC0642-treated mESCs (see Materials and Methods). In this analysis, regions with similar H3K9me2 in WT mESCs were compared. The “early” class showed a higher H3K9me2 level after UNC0642 treatment (Fig. 3D, Additional file 1: Fig. S3B) and overlapped more frequently with G9a/GLP-independent H3K9me2 and SETDB1-dependent H3K9me2 than the “middle” and “late” classes (Fig. 3E, F). In addition, interestingly, H3K9me2 recovery after UNC0642 removal was not induced well in *Setdb1* KO mESCs and TKO iMEFs (Additional file 1: Fig. S3C, D). These data support the idea that G9a/GLP-independent H3K9me2 helps *de novo* H3K9me2 domain formation mediated by G9a/GLP.

**Fig. 3.**
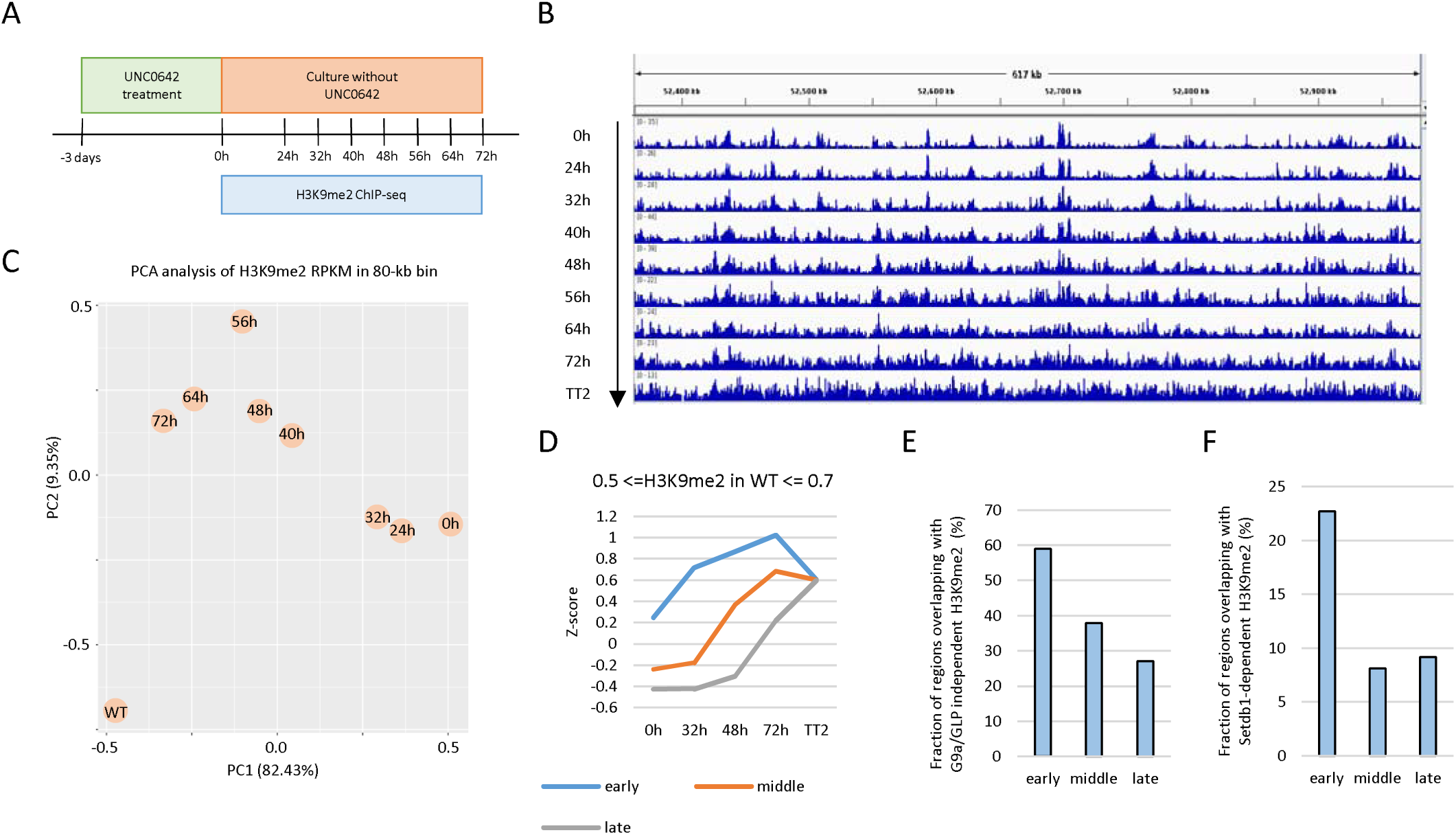
Recovery of H3K9me2 after the UNC0642 treatment A. An experimental design to investigate H3K9me2 recovery. mESCs were treated with UNC0642 for 3 days, and then H3K9me2 ChIP-seq was performed at each time point after the UNC0642 removal. B. A representative view of H3K9me2 ChIP-seq data during H3K9me2 recovery. C. A principal component analysis of H3K9me2 ChIP-seq data during H3K9me2 recovery. D. Distinct kinetics of H3K9me2 recovery from UNC0642 treatment between three classes of sequences. Each 10-kb region throughout the genome was classified to three types named “early”, “middle”, and “late” based on the timing of H3K9me2 recovery. Z-scaling was performed on H3K9me2 RPKM of 10-kb bins in each time point. Y-axis represents the average Z-scores of each class. Only regions with Z-scores between 0.5 and 0.7 in WT mESCs were used for this analysis to match the final H3K9me2 levels among three classes. The regions that show faster H3K9me2 recovery tend to harbor higher H3K9me2 after UNC0642 treatment. E. Overlap between G9a/GLP-independent H3K9me2 domains and each class. “Early” class shows the most frequent overlap with G9a/GLP-independent H3K9me2 regions among three classes. Only regions with Z-scores between 0.5 and 0.7 in WT mESCs were used for this figure. F. Overlap between *Setdb1*-dependent H3K9me2 domains and each class. Only regions with Z-scores between 0.5 and 0.7 in WT mESCs were used for this figure.

### G9a/GLP-independent H3K9me2 is involved in transcriptional repression

To further address the function of G9a/GLP-independent H3K9me2, we investigated whether G9a/GLP-independent H3K9me2, mainly catalyzed by SETDB1, regulates transcription, and performed RNA-seq analysis in the WT and *Setdb1* KO mESCs treated with or without UNC0642. We identified 161, 93, and 75 upregulated genes in UNC0642-treated *Setdb1* KO, *Setdb1* KO, and UNC0642-treated mESCs, respectively (Additional file 1: Fig. S4A). Fifty-four of the 161 upregulated genes in UNC0642-treated *Setdb1* KO mESCs were identified in neither *Setdb1* KO without UNC0642 treatment nor UNC0642-treated WT mESCs, which suggests a redundant function of SETDB1 and G9a/GLP in gene repression (Additional file 1: Fig. S4A). G9a/GLP mainly repressed genes in the B compartments, while SETDB1 repressed genes in both the A and B compartments (Additional file 1: Fig. S4B). Most upregulated genes both in A and B compartments in UNC0642-treated WT mESCs were derepressed in *Setdb1* KO mESCs (17 of 18 in the A compartments and 56 of 57 in the B compartments, FC>2) (Additional file 1: Fig. S4C). In contrast, upregulated genes in the B compartments in *Setdb1* KO mESC was more derepressed in UNC0642 treated mESCs than genes in the A compartments (35 of 56 in the A compartments (62.5%) and 32 of 36 in the B compartments (88.9%), FC>2) (Additional file 1: Fig. S4D). Furthermore, the additive upregulation effect that was observed upon UNC0642 treatment in *Setdb1* KO mESCs was also higher in genes in the B compartments than in the A compartments (Additional file 1: Fig. S4E). Therefore, genes repressed by SETDB1 in the B compartments are co-regulated by G9a/GLP; G9a/GLP-mediated H3K9me2 is more important for genes repressed by SETDB1 in the B compartments than in the A compartments.

To further investigate the association between G9a/GLP-independent H3K9me2 and gene expression, we performed an integrative RNA-seq, H3K9me2 ChIP-seq, and H3K9me3 ChIP-seq analysis. G9a/GLP-independent H3K9me2 was frequently found within 5-kb from the transcriptional start sites (TSS) of upregulated genes in each condition, while H3K9me3 was found less frequently (Fig. 4A). Both H3K9me3 and G9a/GLP-independent H3K9me2 were enriched around the upregulated genes in *Setdb1* KO mESCs and these H3K9me3 and H3K9me2 were dependent on SETDB1 (Fig. 4B, C). In contrast, H3K9me3 around the upregulated genes in the UNC0642-treated mESCs was not dependent on SETDB1, and G9a/GLP-independent H3K9me2 decreased in the *Setdb1* KO mESCs (Fig. 4B, C). Therefore, H3K9me2 correlated with gene repression to a greater extent than H3K9me3. To investigate more clearly whether SETDB1-dependent H3K9me2 functions as a transcriptional repressor, the expression of genes with SETDB1-dependent H3K9me2 but without H3K9me3 around TSS were analyzed (280 genes). Although these genes were not marked with H3K9me3, they were upregulated in the *Setdb1* KO mESCs (Fig. 4D, E). The upregulation of the three selected genes was validated using qRT-PCR (Fig. 4F). These data support the idea that SETDB1-dependent H3K9me2 has a role in transcriptional repression and is not just an intermediate of H3K9me3.

**Fig. 4.**
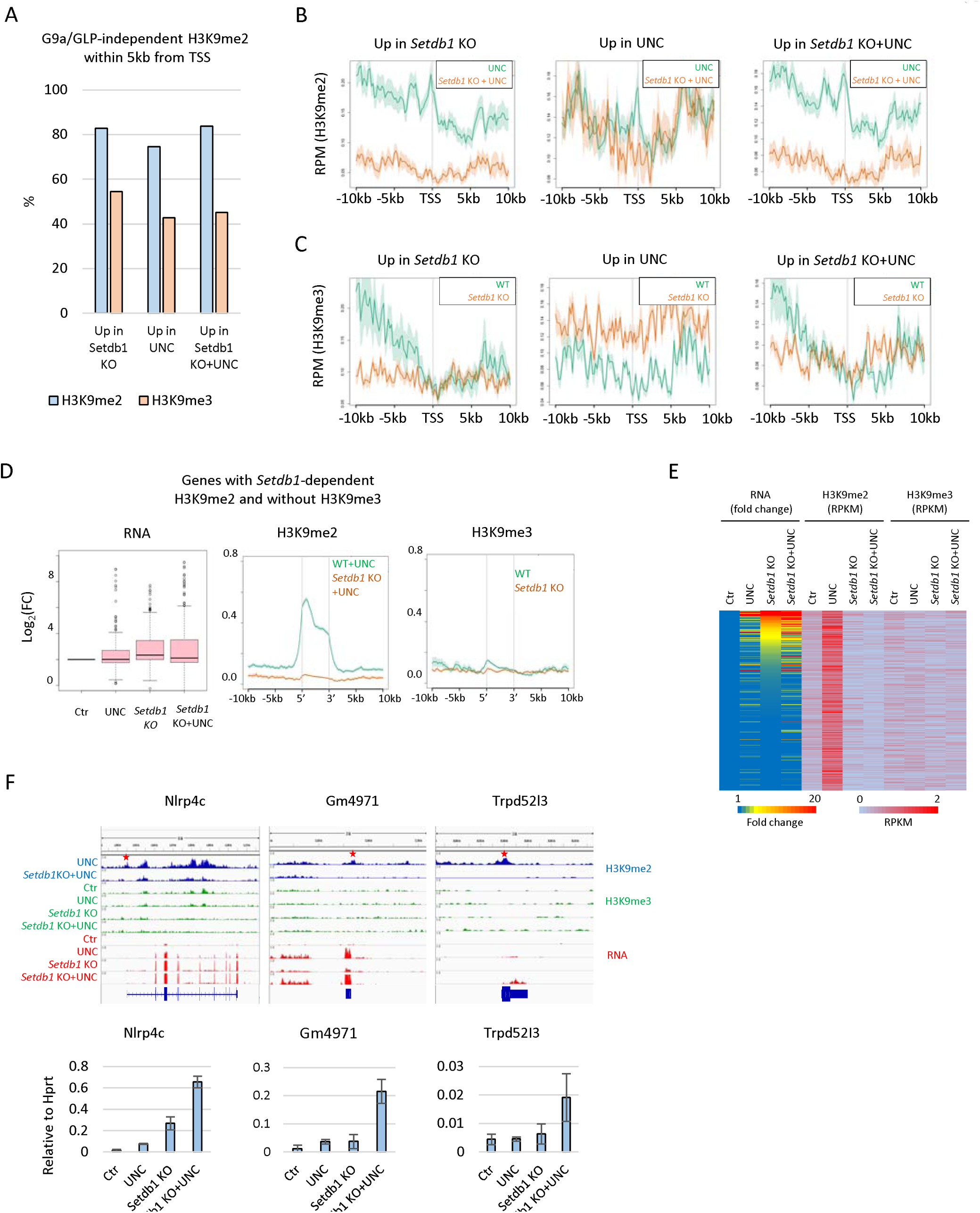
Function of G9a/GLP-independent H3K9me2 in transcriptional regulation A. Overlap of upregulated genes with G9a/GLP-independent H3K9me2 or H3K9me3 domains. The graph shows the fractions of upregulated genes in each condition that overlap with G9a/GLP-independent H3K9me2 domains or H3K9me3 domains within 5kb of TSS. B. Enrichment of H3K9me2 in WT or *Setdb1* mESCs treated with UNC0642 around upregulated genes in each condition. G9a/GLP-independent H3K9me2 is enriched from upstream of TSS to gene body of upregulated genes identified in *Setdb1* KO mESCs treated with or without UNC0642. These H3K9me2 is largely dependent on *Setdb1*. C. Enrichment of H3K9me3 in WT or *Setdb1* mESCs treated with UNC0642 around upregulated genes in each condition. Different from G9a/GLP-independent H3K9me2, H3K9me3 is mainly enriched upstream of upregulated genes identified in *Setdb1* KO mESCs treated with or without UNC0642. D. Expression and H3K9 methylation profiles of genes with Setdb1-dependent H3K9me2 and without H3K9me3. Genes that satisfy the following criteria are extracted: 1. H3K9me2 domains are located within 0.5 kb of TSS in UNC0642-treated mESCs, but not in UNC0642-treated *Setdb1* KO mESCs. 2. H3K9me3 domains are not located within 0.5 kb of TSS in any condition. 3. H3K9me2 RPKM in UNC0642-treated mESCs around 0.5 kb of TSS is more than 1, and that in UNC0642-treated *Setdb1* KO mESCs is less than 1. 4. H3K9me3 RPKM around 0.5 kb of TSS is less than 1 in any condition. The left panel is a boxplot of log2 fold change of expression of genes satisfying the above criteria. Middle figure shows the enrichment of H3K9me2 around the TSS in UNC0642-treated WT or *Setdb1* KO mESCs. The right figure shows the enrichment of H3K9me3 around the TSS in WT or *Setdb1* KO mESCs. E. Heatmap of fold change of expression, H3K9me2 RPKM, and H3K9me3 RPKM in genes extracted in Fig. 4D. H3K9me2 and H3K9me3 RPKM were calculated around 0.5 kb of the TSS. F. qRT-PCR of representative genes extracted in Fig. 4D. Top panel shows H3K9me2, H3K9me3, and RNA-seq profiles of Nlrp4c, Gm4971, and Trpd52I3. Setdb1-dependent H3K9me2 is present around TSS of these genes, but H3K9me3 is absent in all conditions. Star mark represents Setdb1-dependent H3K9me2 around TSS. Bottom graph shows relative expression of these three genes in each condition (N = 3).

### Correlation of decreased H3K9me2 with reorganization of the active compartment setting

Lastly, to clarify whether H3K9me2 has a role in A/B compartment formation, we performed Hi-C analysis in each H3K9 MTase deficient mESC treated with or without UNC0642 and calculated the compartment scores in bins of 100-kb. The Hi-C analysis showed an overall maintenance of nuclear compartment patterns in all the samples analyzed (Fig. 5A), and more than 94% of the compartment profiles were conserved in all the samples (Fig. 5B). Moreover, the overall compartment scores did not largely differ between samples (Additional file 1: Fig. S5A). To elucidate the effect of the H3K9me2 changes on the nuclear compartments more specifically, we compared the changes in H3K9me2 in the UNC0642-treated mESCs to the changes in the compartment scores. This comparison clearly showed that the decreased H3K9me2 upon UNC0642 treatment correlated with the increased compartment score (Fig. 5C). The further decrease in H3K9me2 in the UNC0642-treated *Setdb1* KO mESCs also correlated with the increased compartment scores (Fig. 5D). In addition to the H3K9me2 changes, the increased gene expression in the UNC0642-treated *Setdb1* KO mESCs correlated with increased the compartment scores (Fig. 5E). Such increased compartment scores were observed in both the A and B compartments, and B-to-A compartment changes were also observed in six genes (Fig. 5F). Therefore, decreased H3K9me2 correlated with the re-location of the target genes to more active compartments, accompanied by transcriptional activation.

**Fig. 5.**
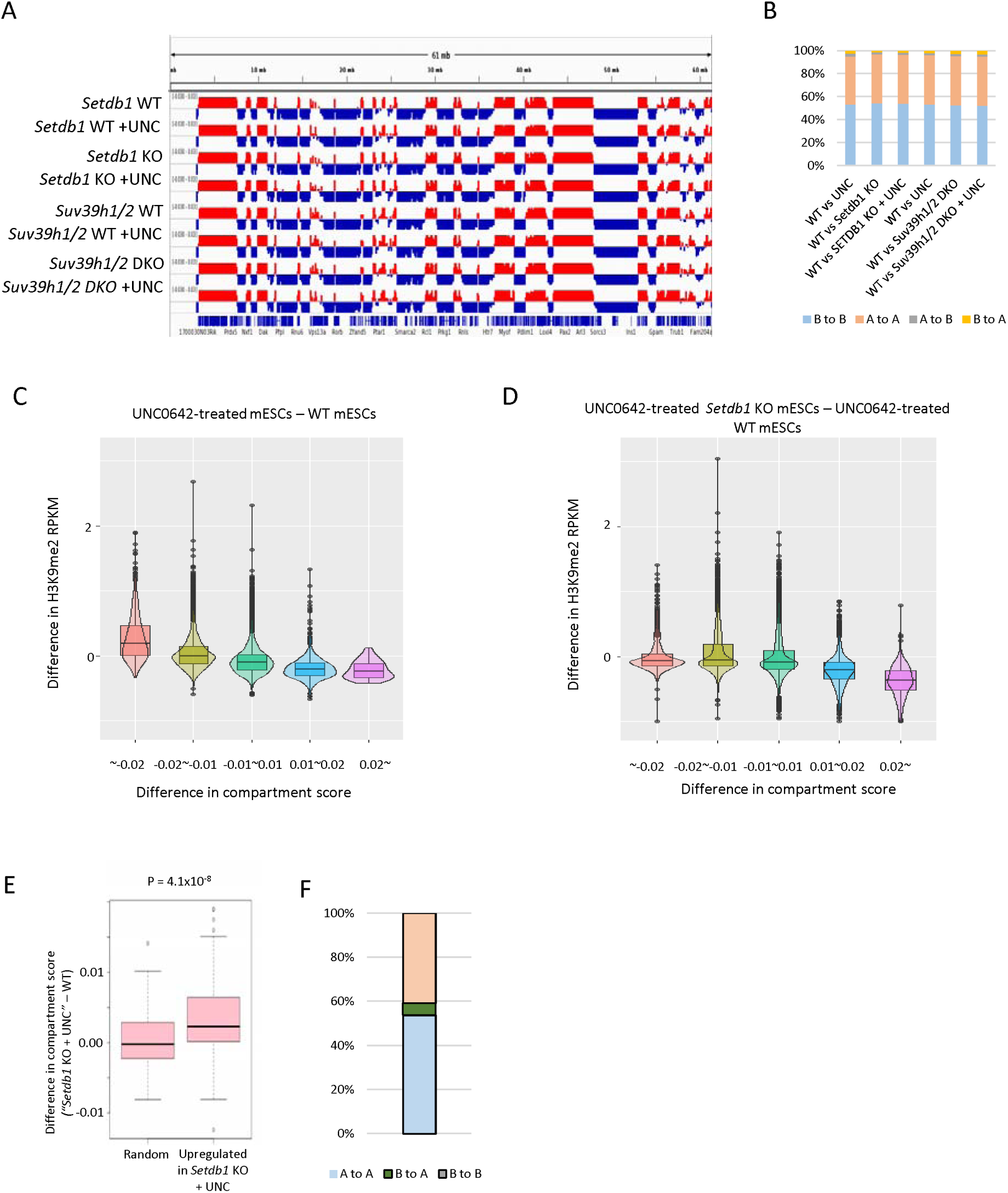
Correlation of decreased H3K9me2 and movement toward more active compartment A. A representative view of compartment score in each 100-kb bin in mESCs. The regions colored by red and blue represent the A and B compartments, respectively. B. Compartment change in each condition. More than 94% of compartments are maintained in all conditions. C. A comparison between change in H3K9me2 and compartment scores in UNC0642-treated mESCs. Each 100-kb region was classified by compartment score changes in UNC0642-treated mESCs. Decreased H3K9me2 in UNC0642-treated mESCs is correlated with an increased compartment score. D. A comparison between changes in H3K9me2 and compartment scores in UNC0642-treated *Setdb1* KO mESCs. Each 100-kb region was classified by compartment score change between UNC0642-treated *Setdb1* KO mESCs and UNC0642-treated mESCs. Decreased H3K9me2 in UNC0642-treated *Setdb1* KO mESCs is correlated with an increased compartment score. E. Change in compartment score around upregulated genes UNC0642-treated *Setdb1* KO mESCs and randomly selected genes. Increased compartment score is observed in upregulated genes in UNC0642-treated *Setdb1* KO mESCs. F. Pattern of compartment changes in genes derepressed in UNC0642-treated mESCs. Only genes of which compartment scores are increased were used in this analysis. Six genes of them showed compartment change from the B compartments to the A compartments.

Overall, this study demonstrated that H3K9me2 is regulated by different sets of H3K9 MTases between the A and B compartments and that G9a/GLP-independent H3K9me2 has a role in efficient H3K9me2 domain formation, transcriptional repression, and 3D genome organization (Fig. 6).

**Fig. 6.**
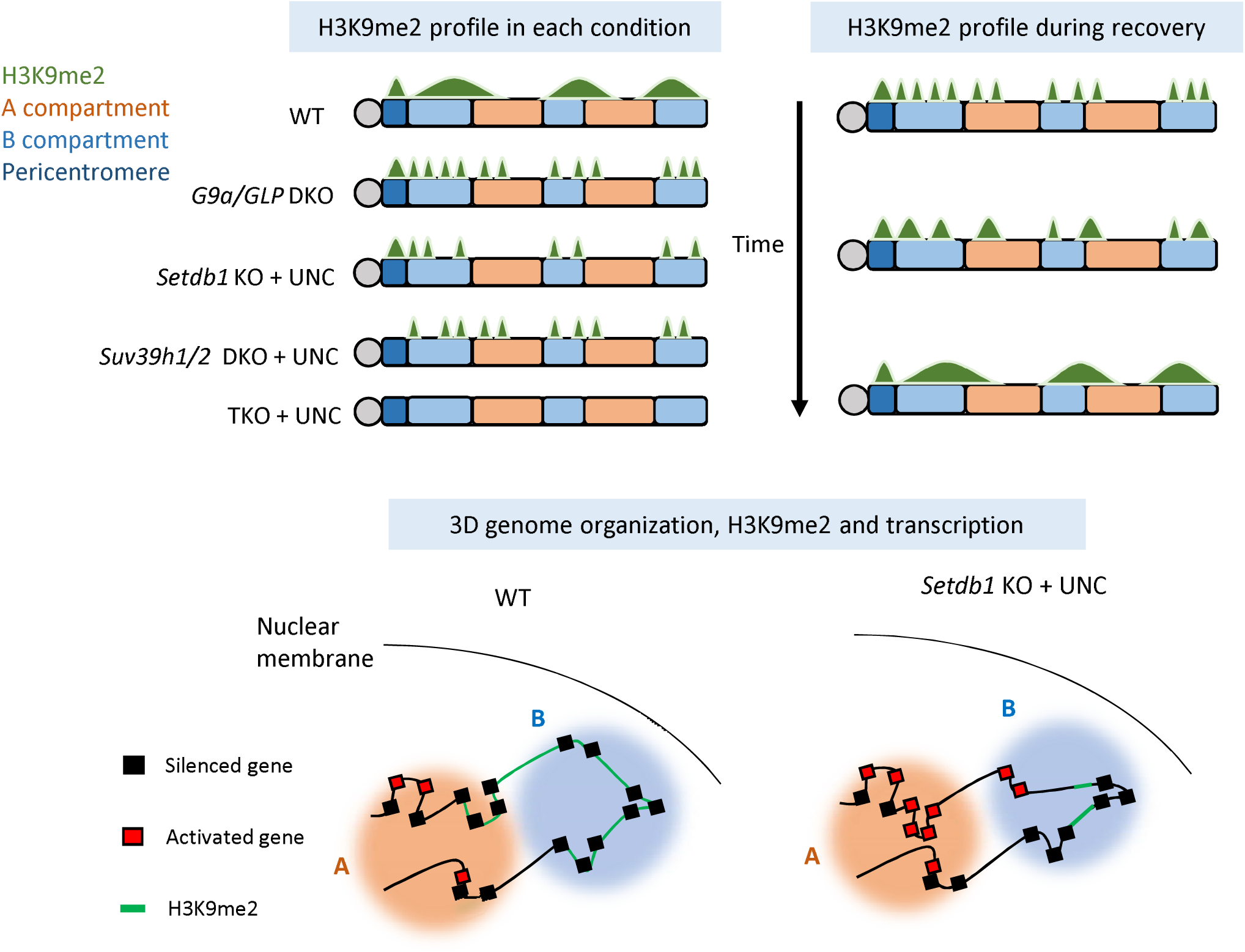
Summary of this study Although H3K9me2 (shown in green) is largely mediated by G9a/GLP in mESCs, some residual H3K9me2 is observed in *G9a/GLP* DKO mESCs. *Setdb1* and *Suv39h1/2* mediate G9a/GLP-independent H3K9me2 in a compartment-dependent manner. *Setdb1* and *Suv39h1/2* are essential for G9a/GLP-independent H3K9me2 in the A compartments (red) and in pericentromeric satellite repeats (dark blue), respectively, while H3K9me2 in the B compartments (light blue) is mediated by both *Setdb1* and *Suv39h1/2* redundantly (Fig. 6 upper left). During H3K9me2 domain establishment after recovery from an UNC0642 treatment, H3K9me2 is recovered efficiently in regions with high G9a/GLP-independent H3K9me2. Thus, G9a/GLP-independent H3K9me2 might facilitate the efficient establishment of H3K9me2 domains (Fig. 6 upper right). Finally, we compared H3K9me2, transcription and 3D genomes, and found that the reduction of H3K9me2 resulted in a movement toward more active compartment, accompanied by transcriptional activation (Fig. 6 bottom). Thus, G9a/GLP-independent H3K9me2 functions in H3K9me2 domain establishment, transcriptional repression and 3D genome organization.

## Discussion

This study investigated the role of five different H3K9 MTases in H3K9me2 domain formation and the function of G9a/GLP-independent H3K9me2 in transcriptional repression and 3D genome organization. Although many studies have implicated the presence of G9a/GLP-independent H3K9me2 in mESCs and iMEFs [6, 14, 15], the mechanism and function of G9a/GLP-independent H3K9me2 are unknown. G9a/GLP-independent H3K9me2 is enriched in pericentromeric satellite repeats in mESCs [6, 14] and observed in H3K9me3-marked regions in mESCs and late-replicating domains in MEFs [15]. Therefore, it has been predicted that G9a/GLP-independent H3K9me2 is mediated by other H3K9 MTases, such as SETDB1 and SUV39H1/2. We formally demonstrated that G9a/GLP-independent H3K9me2 is mediated by SETDB1 and SUV39H1/2. Surprisingly, the contribution of SETDB1 and SUV39H1/2 to H3K9me2 differed in each compartment. SETDB1 is essential for G9a/GLP-independent H3K9me2 in the A compartment, while both SETDB1 and SUV39H1/2 mediate this process in the B compartment (Fig. 6 upper left), and it remains unknown why SUV39H1/2 function is restricted only to the B compartment. A mixture of heterochromatin protein 1 (HP1)-SUV39H1-TRIM28 complexes, H3K9me2, and 3-marked nucleosomal arrays undergoes phase separation *in vitro* [29]. HP1 and SUV39H1 are enriched in DAPI-dense regions and form droplet-like structures in cultured cells [30]. Thus, one possible explanation for SUV39H1 restriction to the B compartment is phase separation and the other possible reason could be the genomic distribution of SUV39H1/2-target regions. SUV39H1/2 represses LINE1 retrotransposon, and LINE1 is enriched in AT-rich isochores and gene-poor regions, which is consistent with the features of the B compartment [31]. Thus, the enrichment of LINE1 in B compartments might restrict SUV39H1/2 function in this location.

Heterochromatin can spread from the nucleation sites. G9a and Glp can bind to H3K9me1 and H3K9me2 via their ankyrin repeat domain [27], and Glp might premethylate nucleosomes and further methylate the premethylated neighboring nucleosomes [28]. We demonstrated that H3K9me2 reduction upon UNC0642 treatment recovered more efficiently after inhibitor removal in the regions with high G9a/GLP-independent H3K9me2 (Fig. 6 upper right). In addition, SETDB1 depletion resulted in no H3K9me2 recovery in mESCs (Additional file 1: Fig. S3C). These results imply that G9a/GLP-independent H3K9me2 helps efficient H3K9me2 establishment or spreading. The HP1-CAF1-SETDB1 complex monomethylates K9 on non-nucleosomal histone H3 [32], and H3K9me1 is imposed during translation by SETDB1 [33]. Thus, H3K9me1 on non-nucleosomal histone H3 mediated by SETDB1 may also provide a scaffold for G9a/GLP during H3K9me2 domain establishment. As H3K9me2 domains are established in intergenic regions in mouse oocytes by G9a, and the global H3K9me2 level is decreased in *Setdb1* KO oocytes [34, 35], SETDB1-mediated G9a/GLP-independent H3K9me2 and/or SETDB1-mediated non-nucleosomal H3K9me1 may be important for H3K9me2 domain establishment during gametogenesis.

In addition to H3K9me2 spreading, our transcriptome analysis demonstrated that G9a/GLP-independent H3K9me2 plays a role in transcriptional repression. Some SETDB1 target genes harbor SETDB1-dependent H3K9me2 but not H3K9me3 (Fig. 4D-F). It is unknown why histone methylation patterns differ among SETDB1 target genes. As SETDB1 catalytic activity is regulated by ATF7IP and the monoubiquitination of lysine 867 of the hSETDB1, monoubiquitination of SETDB1 or ATF7IP enrichment at the target genomic regions might affect the catalytic activity of SETDB1 [4, 36].

Our Hi-C analysis showed that the overall nuclear compartment patterns were maintained in all the analyzed samples. However, H3K9me2 decreased upon UNC0642 treatment in mESCs, which correlated well with increased compartment scores, and the further decrease in H3K9me2 in UNC0642-treated *Setdb1* KO mESCs also correlated with the further increase in the compartment scores (Fig. 5C, D). Therefore, we propose that H3K9me2 contributes to heterochromatin compartment formation as one of the key players of determinant mechanisms (Figure 6 bottom). Because *Setdb1* KO mESCs treated with UNC0642 still possess H3K9me2 in the B compartment (Fig. 6 upper left) but H3K9me2 and H3K9me3 are almost depleted in *Setdb1, Suv39h1/2* TKO mESCs, and iMEFs treated with UNC0642, it will be of interest to analyze genome compartmentalization in this cell line in order to fully dissect the role of H3K9 methylation in this process.

## Methods

### Cell culture

mESCs were maintained in Dulbecco’s modified Eagle’s medium (Sigma) containing 10% Knockout SR (Invitrogen), 1% fetal bovine serum (Biowest), MEM Non-Essential Amino Acids Solution (Gibco), 0.1% LIF and 2-Mercaptoethanol (Nacalai tesque) described as ES medium hereafter). Mouse embryonic fibroblasts were maintained in Dulbecco’s modified Eagle’s medium (Nacalai tesque) containing 10% fetal bovine serum (Biosera), MEM Non-Essential medium and 2-Mercaptoethanol (Nacalai tesque). To inhibit G9a/GLP catalytic activity, mESCs and iMEFs were cultured for three days with 2 μM UNC0642. For conditional *Setdb1* KO, *Setdb1^fl/fl^* mESCs were cultured for five days with 800 nM 4-OHT, while *Suv39h1/2 DKO/Setdb1 ^fl/fl^* mESCs were cultured for seven days with 800 nM 4-OHT.

### Establishment of *Setdb1*, *Suv39h1/2* TKO cells

*Setdb1* (conditional), *Suv39h1/2* TKO mESCs: *Setdb1* conditional KO mESC, #33-6 [8]was transfected with the following four gRNAs: 1. *Suv39h1* exon 3 upstream in PL-CRISPR.EFS.tRFP, 2. *Suv39h2* exon 4 downstream in pKLV2-U6.gRNA (Bbs1)-PGK.puro-BFP, 3. *Suv39h2* exon 3 in PL-CRISPR.EFS.tRFP, 4. *Suv39h2* exon 4 in pKLV2-U6.gRNA (Bbs1)-PGK.puro-BFP. 3 days after the transfection, BFP and RFP double-positive cells were sorted by flow cytometry. From the sorted cells, *Suv39h1/2* DKO mES clone #4 was identified using PCR and western blotting. Partial deletion of *Suv39h1/2* SET domains were validated by RNA-seq analysis (Fig. S1J).

*Setdb1*, *Suv39h1/2* TKO iMEFs: *Setdb1* KO iMEFs [24]were transfected with *Suv39h1* exon 4 gRNA in pKLV2-U6.gRNA(Bbs1)-PGK.puro-BFP and *Suv39h2* exon 4 upstream and downstream gRNAs in pX330-BB and selected with puromycin. From the puromycin resistant cells, *Setdb1, Suv39h1* DKO iMEF clone #27 was identified by Western blot analysis. Then, *Suv39h1* KO clone #27 was further transfected with *Suv39h2* exon 3 gRNA in pL-CRISPR.EFS.tRFP and pKLV2-U6.gRNA(Bbs1)-PGK.puro-BFP and selected with Puromycin again. From the puromycin resistant cells, *Setdb1*, *Suv39h1/2* TKO KO cell clone #27-34 was identified by Western blot analysis. CRISPR-Cas9 mediated *Suv39h1* and 2 gene mutations were confirmed by DNA sequencing analysis (Fig. S2C).

### Native ChIP

At least 2×10^5^ cells were lysed in 50 ul Buffer 1 (60mM KCl, 15mM NaCl, 5mM MgCl2, 0.1 mM EGTA, 15mM Tris-HCl (pH7.5), 0.3M Sucrose, 0.5mM DTT, and protease inhibitors), then were added 50 ul Buffer2 (Buffer 1 + 1% NP40) to the sample. After the incubation for 10 min on ice, the sample was added 800 ul Buffer 3 (60mM KCl, 15mM NaCl, 5mM MgCl2, 0.1mM EGTA, 15mM Tris-HCl (pH7.5), 1.2M Sucrose, 0.5mM DTT, and protease inhibitors). After centrifugation at 9,000 rpm for 10min at 4°C, the sample was added 100ul MNase buffer (0.32M Sucrose, 50mM Tris-HCl (pH7.5), 4mM MgCl2, 1mM CaCl2, and PMSF) followed by incubation for 20min at 37°C with 0.3U MNase (TAKARA). After the incubation, the reaction was stopped by 10 ul 0.5M EDTA. After centrifugation at 15,000 rpm for 10min at 4°C, the supernatant was added 900 ul Incubation buffer (50mM NaCl, 20mM Tris-HCl (pH7.5), 5mM EDTA, 0.01% NP40, ad PMSF). 10% of the sample was used for input, and the remains were proceeded the following procedure. The sample was mixed with the antibody-beads complex which was formed by incubation of 20ul protein A/G beads with antibody in Incubation buffer on ice for 1h. After rotating for overnight at 4°C, the complex was washed by 500ul Wash buffer A (75mM NaCl, 50mM Tris-HCl (pH7.5), 10mM EDTA, and 0.01% NP40), 500ul Wash buffer B (100mM NaCl, 50mM Tris-HCl (pH7.5), 10mM EDTA, and 0.01% NP40), and 500ul Wash buffer C (175mM, 50mM Tris-HCl (pH7.5), 10mM EDTA, and 0.01% NP40). The ChIP DNA was eluted by RNase and Protease treatment followed by DNA purification using PCR purification kit (QIAGEN).

### Western blotting

Briefly, cells were suspended in RIPA buffer and sonicated. The extract was incubated for 30 minutes on ice, and then incubated at 95°C for 5 minutes. The extract was loaded and run on SDS-PAGE gel as standard protocols. For histone proteins, intensity was analyzed by Odyssey Clx (LI-COR).

### Mass spectrometry analysis for H3K9 methylation

Histones were prepared by the acid-extraction method as previously described [6]. The samples were subjected to SDS-PAGE and stained with Coomassie blue. The protein bands corresponding to histone H3 were excised from the gel and were digested with *Achromobacter* protease I (Lys-C) in gel. The digests were performed nano-liquid chromatography - tandem mass spectrometry using EASY-nLC 1200 liquid chromatograph (Thermo Fisher Scientific) connected to Q-Exactive HFX mass spectrometer equipped with nanospray ion source (Thermo Fisher Scientific). Peptides containing the digests were separated with a linear gradient of 0-100%B buffer in A buffer (A:0.1% formic acid, B:80%acetonitrile/0.1%formic acid) for 20min with a reversed-phase column (NTCC analytical column, C18, φ75 μm ×150 mm, 3 μm; Nikkyo Technos, Japan). MS and MS/MS data were acquired using a data dependent TOP 10 method. Protein quantification were performed with Proteome Discoverer 2.4. (Thermo Fisher Scientific) using a sequence database search node as a MASCOT program 2.7 (Matrix Science) with following parameters, Database: Histone 130611 (66 sequences; 15002 residues), Enzyme: Lys-C/P, Variable modifications: Acetyl (Protein N-term),Oxidation (M),Gln->pyro-Glu (N-term Q),Acetyl (K),Methyl (K),Trimethyl (K),Dimethyl (K), Mass values: Monoisotopic, Protein Mass: Unrestricted, Peptide Mass Tolerance: ± 15 ppm, Fragment Mass Tolerance: ± 30 mmu, Max Missed Cleavages: 3, Instrument type: ESI-TRAP. Protein methylation rates were obtained peak areas of selected ion chromatograms that are the tetra charged protonated molecules at m/z 381.9706 (methyl), m/z 385.4745 (dimethyl) and m/z 388.9784 (trimethyl) of “QTARKSTGGKAPRK” related peptides (1xAcetyl [K10]; 1xGln->pyro-Glu [N-Term]; 1xmethylations [K5]) using Qual Browser (Thermo xcalibur 4.1.50, Thermo Fisher Scientific) with ±15ppm width.

### RT-qPCR

RNA was isolated by RNeasy Plus Mini Kit (Qiagen) following manufacturer’s instructions. cDNA synthesis was performed with Omniscript RT Kit (Qiagen). qPCR was carried out using Power SYBR Green PCR Master Mix (ThermoFisher SCIENTIFIC) on StepOnePlus™ (ThermoFisher SCIENTIFIC). The signals were normalized relative to *Hprt*. All qPCR data is represented as the mean +/-standard deviation of three biological replicates.

### Preparation of ChIP-seq library

The ChIP DNA was fragmented by Picoruptor (Diagenode) for 10 cycles of 30 seconds on, 30seconds off. Then, ChIP library was constructed by KAPA Hyper Prep Kit (KAPA BIOSYSTEMS) and SeqCap Adapter Kit A (Roche) or SMARTER ThruPLEX DNA-seq kit (TAKARA) and SMARTer DNA Unique Dual Index Kit (TAKARA) according to manufacturer instructions. The concentration of the ChIP-seq library was quantified by KAPA Library quantification kit (KAPA BIOSYSTEMS). ChIP sequencing was performed on a HiSeq 2000 platform (Illumina).

### Preparation of RNA-seq library

500 ng of total RNA was used for RNA-seq library construction. RNA-seq library was constructed by KAPA mRNA Hyper Prep Kit (KAPA BIOSYSTEMS) and SeqCap Adapter Kit (Roche) according to manufacturer instructions. The concentration of the RNA-seq library was quantified by KAPA Library quantification kit (KAPA BIOSYSTEMS). mRNA sequencing was performed on a HiSeq 2000 platform (Illumina).

### Preparation of HiC-seq library

Hi-C experiments were performed as previously described [37–39], based on DpnII enzyme (4-bps cutter) using 2×10^6^ fixed cells. Hi-C libraries were subject to paired-end sequencing (150 base pair (bp) read length) using HiSeq X Ten.

### Hi-C data processing and A/B compartment calculation

Hi-C data processing was done by using Docker for 4DN Hi-C pipeline (v43, https://github.com/4dn-dcic/docker-4dn-hic). The pipeline includes alignment (using the mouse genome, mm10) and filtering steps. After filtering valid Hi-C alignments,.*hic* format Hi-C matrix files were generated by Juicer Tools [40] using the reads with MAPQ>10. The A/B compartment (compartment score) profiles (in 100-kb bins) in each chromosome (without sex chromosome) were calculated from.*hic* format Hi-C matrix files (intrachromosomal KR normalized Hi-C maps) by Juicer Tools [40] as previously described[41].

### ChIP-seq analysis

#### ·Mapping and domain identification

Adaptor sequences in reads were trimmed using Trim Galore version 0.3.7 (http://www.bioinformatics.babraham.ac.uk/projects/trim_galore/). Then trimmed reads were aligned to the mm10 genome build using bowtie version 0.12.7 [42] with default parameters. Duplicated reads were removed using samtools version 0.1.18 [43]. H3K9me2 domains were identified by *Hiddendomains* with some modifications: The option of Bin size and max.read.count was 2,000 bp and 150, respectively. Read number in each bin was normalized by the following formula to adjust difference in read number among samples. Normalized read number = read number x 30,000,000 / total read number. For the hidden Markov parameter to determine enriched and depleted states, average parameter among chromosomes rather than parameter calculated by each chromosome was used, because very few domains were identified in some chromosomes. Loss of H3K9me2 domains was identified using getDifferentialPeaks in Homer (fold change >= 4 and P-value <= 0.0001). Conserved H3K9me2 domains ware the domains where was identified also in a sample and were not identified as lost domains.

#### ·Classification of regions by the timing of H3K9me2 recovered

RPKM of each 10-kb bin was calculated and the RPKM was converted to z-score. The 10-kb bin where Z-score increased by 0.3 or more at 32h, 48h, and 72h from 0h was annotated as early, middle, and late, respectively.

### RNA-seq analysis

Raw FastQ data were trimmed with Trim Galore (v0.3.7, default parameters) (http://www.bioinformatics.babraham.ac.uk/projects/trim_galore/) and mapped to the mouse GRCm38 genome assembly using TopHat (v2.1.1) [44]. After read mapping, mapped reads were analyzed by TEtranscripts (v1.4.11, default parameters) [45] to calculate gene expression levels and identify DE genes (adj. P-value < 0.05, FC > 10).

### Visualization of NGS data

The Integrative Genomics Viewer (IGV) was used to visualize NGS data. Enrichment of H3K9me2/H3K9me3 enrichment in specified regions was visualized by ngsplot. Scatter plot analysis, principal component analysis boxplot and violin plot analysis were performed by R script.

### Immunofluorescence analysis

2×10^4^ cells were seeded on laminin coated 12 well Chamber (Ibidi) the day before fixation. The cells were fixed with 4% paraformaldehyde for 10 minutes at room temperature, permeabilized with 0.1% Triton X-100 for 10 minutes, blocked with 3% BSA 0.1% Tween20 in PBS and incubated overnight with primary antibodies at 4°C. Anti-mouse IgG conjugated with Alexa Fluor 568 (ThermoFisher SCIENTIFIC) or anti-rabbit IgG conjugated with Alexa Fluor 488 (ThermoFisher SCIENTIFIC) were used as secondary antibodies. The nuclei were counterstained with DAPI, observed under fluorescence microscopy and analyzed Olympus FluoView™ FV3000 (Olympus).

### Antibodies

For western blotting, antibodies specific for histone H3 (07-690, EMD Millipore), H3K9me3 (2F3), H3K9me2 (6D11), *SETDB1* (Cp10377, Cell Applications), Suv39h1 (#8729, CST), *Suv39h2* (LS-116360, LSBio), *G9a* (A8620A) and *Glp* (B0422B) were used as primary antibody. For histone proteins, IRDye 800CW Goat anti-Mouse IgG (926-32210, LI-COR) and IRDye 680RD Goat anti-Rabbit IgG (926-68071, LI-COR) were used as secondary antibody. For non-histone proteins, HRP-linked anti-Rabbit IgG (NA934, GE Healthcare) and HRP-linked anti-Mouse IgG (NA931, GE Healthcare) were used as secondary antibody. For immunofluorescence analysis, H3K9me2 (6D11), H3K9me3 (39161, Active Motif) were used for primary antibody, and Goat anti-Mouse IgG Alexa Fluor 568 (A-11031, Invitrogen) and Goat anti-Rabbit IgG Alexa Fluor 488 (A-11034) were used for secondary antibody. For ChIP analysis, antibodies specific for H3K9me3 (2F3) and H3K9me2 (6D11) were used.

### Oligonucleotides

Oligonucleotide sequences for PCR primers and production of gRNA targeting vectors are listed in Supplementary Table S1 and S2, respectively.

## DATA ACCESS

All reads from the RNA-seq, ChIP-seq and Hi-C experiments in this study have been submitted to Sequence Read Archive (SRA; https://www.ncbi.nlm.nih.gov/sra) under accession number SRPXXXX. Read number of NGS data are listed in Supplementary Table S3.

## Acknowledgement

We thank Thomas Jenuwein (Max Planck Institute) for providing *Suv39h1/2* DKO mES and iMEF cell lines, the staff of the Support Unit for Bio-Material Analysis (BMA) at RIKEN Center for Brain Science (CBS) Research Resources Division (RRD) for DNA sequencing, and NGS library construction with special thanks to K. Ohtawa. We also thank all of the Cellular Memory laboratory members for helpful discussions.

## Author contributions

Conceptualization: YS, KF. Cell culture and western blotting analysis: CS, KF. NGS analysis: KF, AT, HM, HI. Mass spectrometry analysis: TS, ND. Paper writing: YS, KF, IH, TD.

## Ethics approval and consent to participate

Not applicable.

## Consent for publication

Not applicable.

