## Supplemental Tables for "Functional correlation of H3K9me2 and nuclear compartment formation": 200824 H3K9me2 Sup Figure2.pdf

### *Supplementary Materials*

### Fukuda et al. Fig. S1

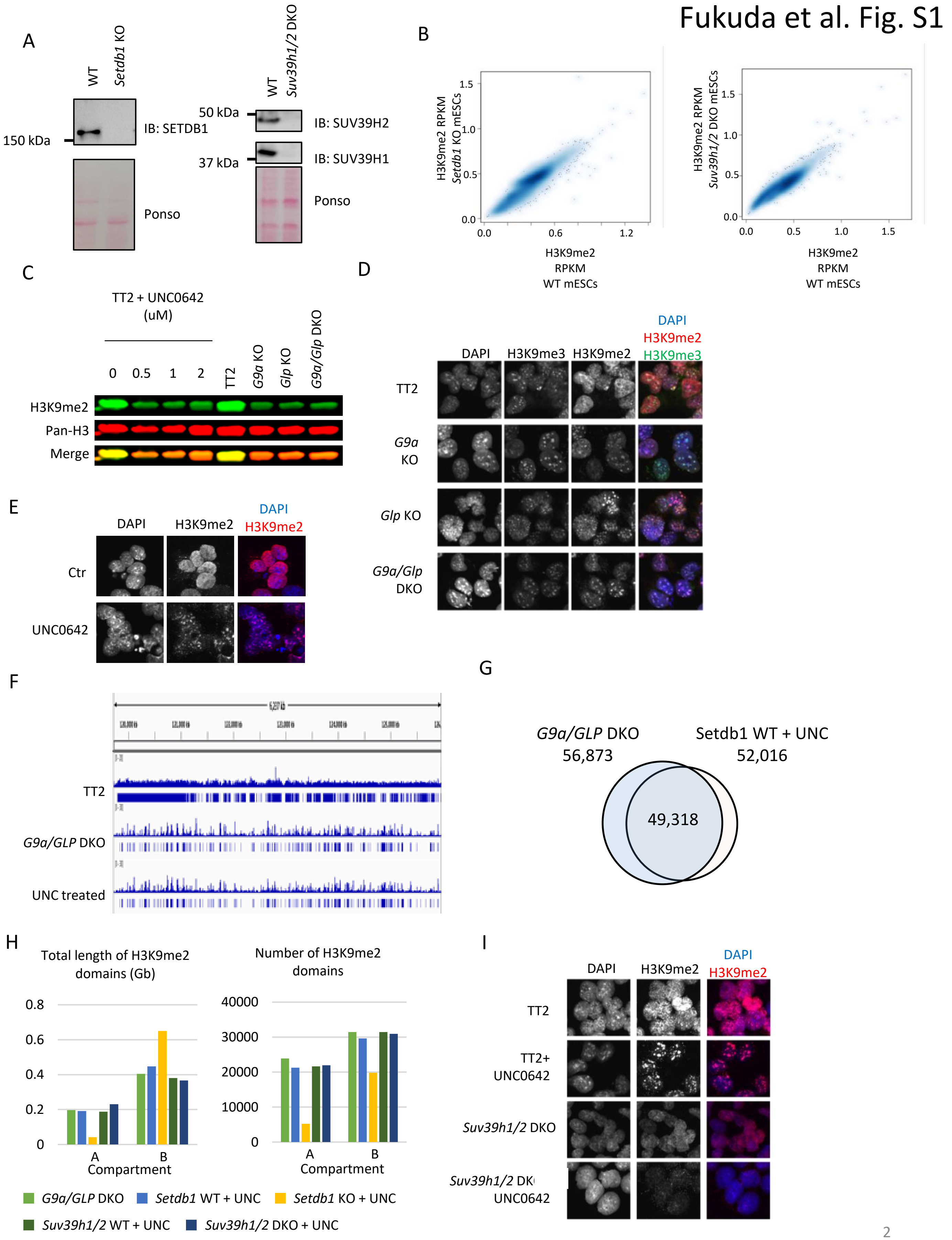

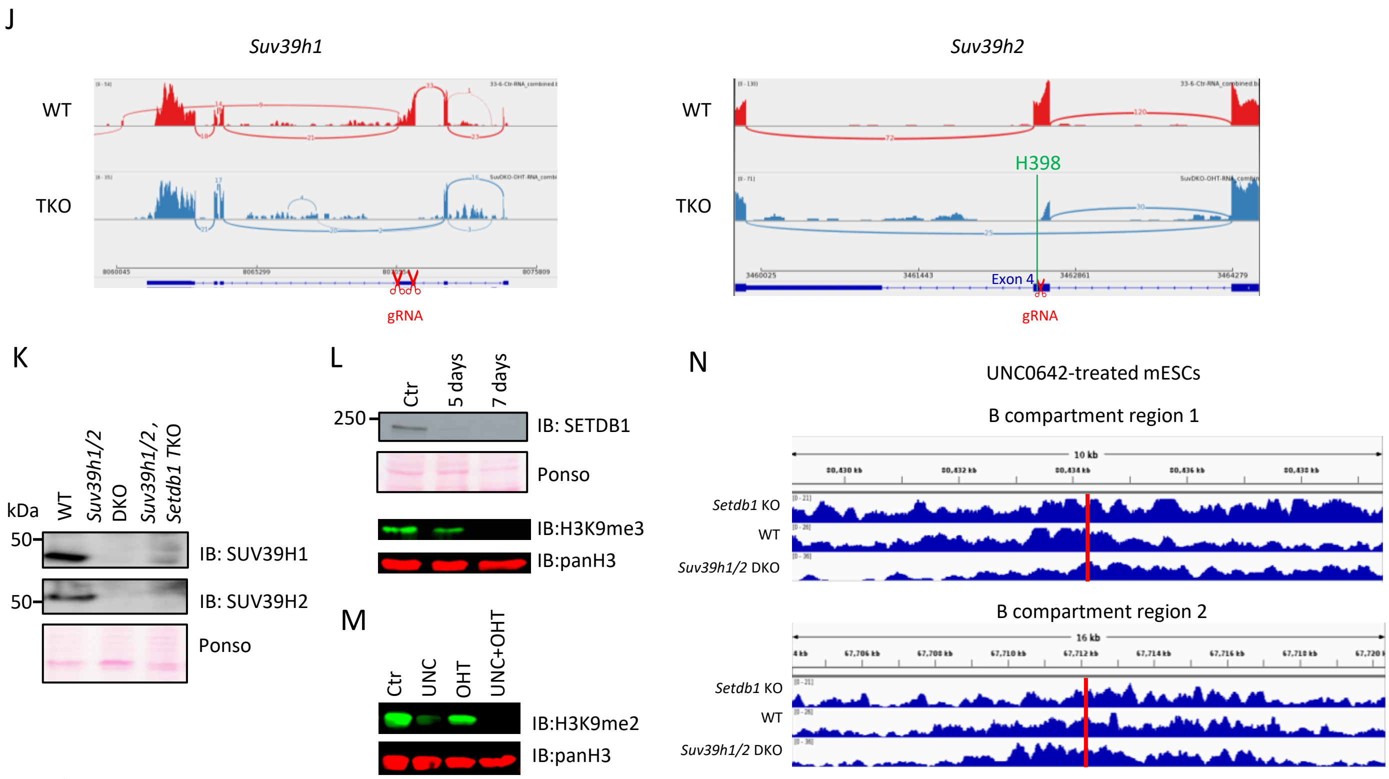

Fig. S1

A. Validation of *Setdb1* KO and *Suv39h1/2* DKO mESCs by western blotting. SETDB1 protein is not detectable five days after 4-OHT treatment in *Setdb1* conditional KO mESCs.

B. Comparison of H3K9me2 RPKM in 80-kb bins between WT and *Setdb1* (left) or *Suv39h1/2* KO (right) mESCs. H3K9me2 profiles are not changed largely in *Setdb1* or *Suv39h1/2* KO mESCs. Darker blue represents higher dot density.

C. Western blotting analysis of H3K9me2 in *G9a/GLP* KO mESCs and UNC0642-treated mESCs. UNC0642 treatment for three days reduces H3K9me2 to levels comparable to *G9a* and/or *GLP* KO mESCs.

D. Immunofluorescence analysis of H3K9me2/3 in *G9a/GLP* KO mESCs. While H3K9me2 is localized in the entire nucleoplasm in WT mESCs, it is enriched in DAPI-dense loci in *G9a/GLP* KO mESCs.

E. Immunofluorescence analysis of H3K9me2 in UNC0642-treated mESCs. Consistent with *G9a/GLP* KO mESCs, H3K9me2 is enriched in DAPI-dense loci in UNC0642-treated mESCs.

F. A representative view of H3K9me2 ChIP-seq data in WT, *G9a/GLP* DKO, and UNC0642-treated mESCs. Blue boxes represent H3K9me2 domains identified by *Hiddendomains*. The size of H3K9me2 domains in *G9a/GLP* or UNC0642-treated mESCs are smaller than those in WT mESCs.

G. An overlap of H3K9me2 domains between *G9a/GLP* DKO mESCs and UNC0642-treated mESCs. 94.8% of H3K9me2 domains in UNC0642-treated mESCs is overlapped with those in *G9a/GLP* DKO mESCs.

H. The length and the number of H3K9me2 domains in *G9a/GLP* DKO or UNC0642-treated mESCs. In UNC0642-treated *Setdb1* KO mESCs, both the total length and the number of H3K9me2 domains in the A compartments are decreased. Immunofluorescence analysis of H3K9me2 in UNC0642-treated *Suv39h1/2* DKO mESCs. H3K9me2 enrichment in DAPI-dense loci observed in UNC0642-treated mESCs is lost in UNC0642-treated *Suv39h1/2* DKO mESCs, suggesting an essential role of SUV39H1/2 for H3K9me2 in pericentromeric satellite repeats.

I. Validation of *Suv39h1* and *2* mutation in TKO mESCs by RNA-seq analysis. Sahimi-plot was generated by Integrative Genome Viewer. Exon 3 of *Suv39h1* is completely deleted in TKO mESCs. *Suv39h1* is X-linked gene and one copy in the TKO mESC (male). H398 of SUV39H2, which is an essential for the methyltransferase activity, is removed from the *Suv39h2* transcripts.

J. Validation of *Suv39h1/2* KO in *Setdb1/Suv39h1/2* TKO mESCs by western blotting.

K. Validation of *Setdb1* KO and H3K9me3 loss by western blotting in TKO mESCs. five days after 4-OHT treatment is not enough to SETDB1 and H3K9me3 depletion. SETDB1 and H3K9me3 are not detectable seven days after 4-OHT treatment.

L. Validation of H3K9me2 loss in *Setdb1/Suv39h1/2* TKO mESCs by western blotting. H3K9me2 is not detectable in UNC0642-treated TKO mESCs 7 days after 4-OHT treatment.

M. Representative B compartment regions in which H3K9me2 remains in both *Setdb1* KO and *Suv39h1/2* DKO mESCs treated with UNC0642. Red bars indicate a region analyzed by ChIP-qPCR in Figs. 1J and 2G.

### Fukuda et al. Fig. S2

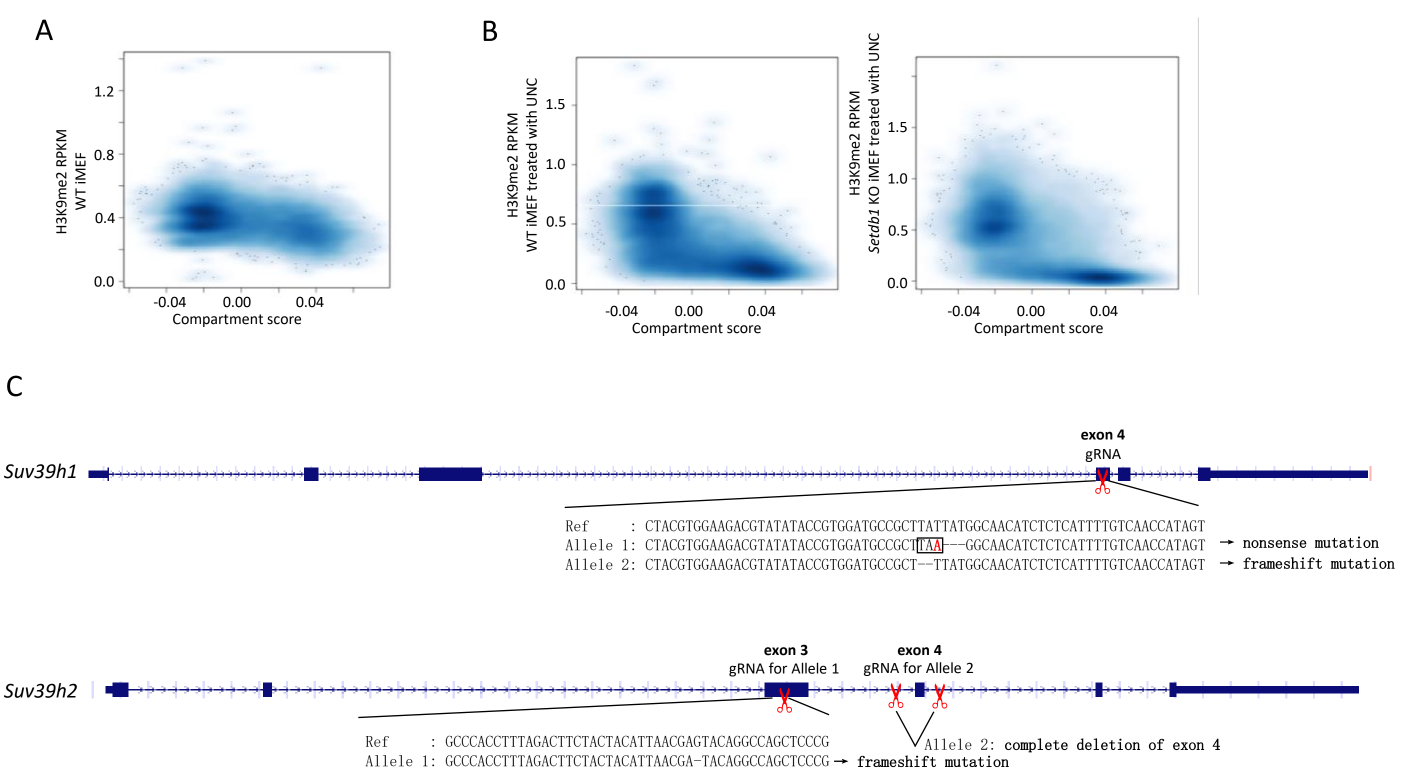

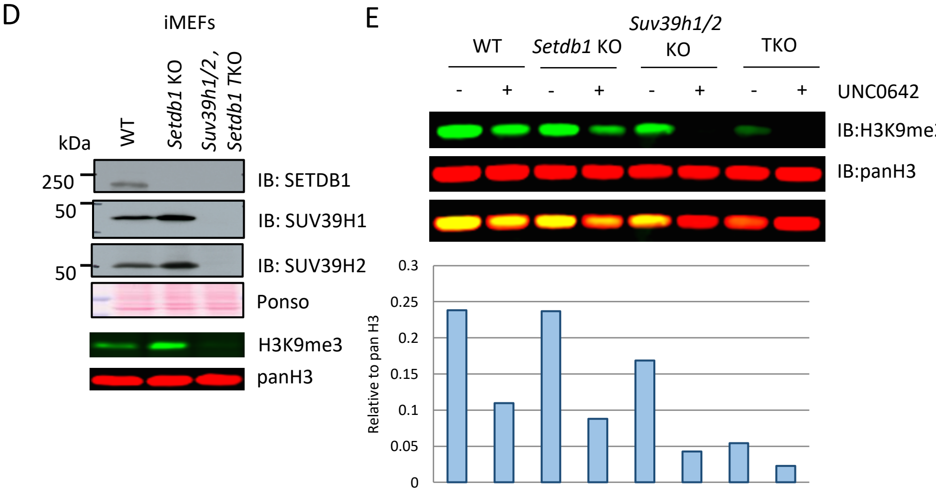

F

| Relative to WT | WT+UNC | TKO | TKO+UNC |
|---|---|---|---|
| H3K9me1 | 44% | 188% | 14% |
| H3K9me2 | 24% | 30% | 1% |
| H3K9me3 | 35% | 1% | 0% |

Fig. S2

A. A scatter plot between compartment score and H3K9me2 RPKM in iMEFs. Compartment score in mESCs was used for the analysis. Each plot represents data from 80-kb bin . Darker blue represents higher dot density.

B. Scatter plots between compartment score and H3K9me2 RPKM in WT (left) and *Setdb1* KO (right) iMEFs treated with UNC0642. Lower H3K9me2 in the A compartments (positive value region) in UNC0642-treated *Setdb1* KO iMEFs than in UNC0642-treated WT iMEFs is observed.

C. Genotype of *Setdb1/Suv39h1/2* TKO iMEFs, which is derived from female mice.

D. Validation of *Setdb1/Suv39h1/2* TKO iMEFs by western blotting. H3K9me3 is not detectable in TKO iMEFs.

E. Western blotting analysis of H3K9me2 in TKO iMEFs. H3K9me2 is not detectable in UNC0642-treated TKO iMEFs. Bar graph at the bottom shows amounts of H3K9me2 relative to pan-H3.

F. A mass spectrometry analysis of H3K9 methylation in iMEFs. iMEFs were treated with $2\mu M$ of UNC0642 for five days. Relative amounts of H3K9 methylation to WT are shown. Both H3K9me2 and H3K9me3 were essentially lost in UNC0642-treated TKO iMEFs.

Fukuda et al. Fig. S3

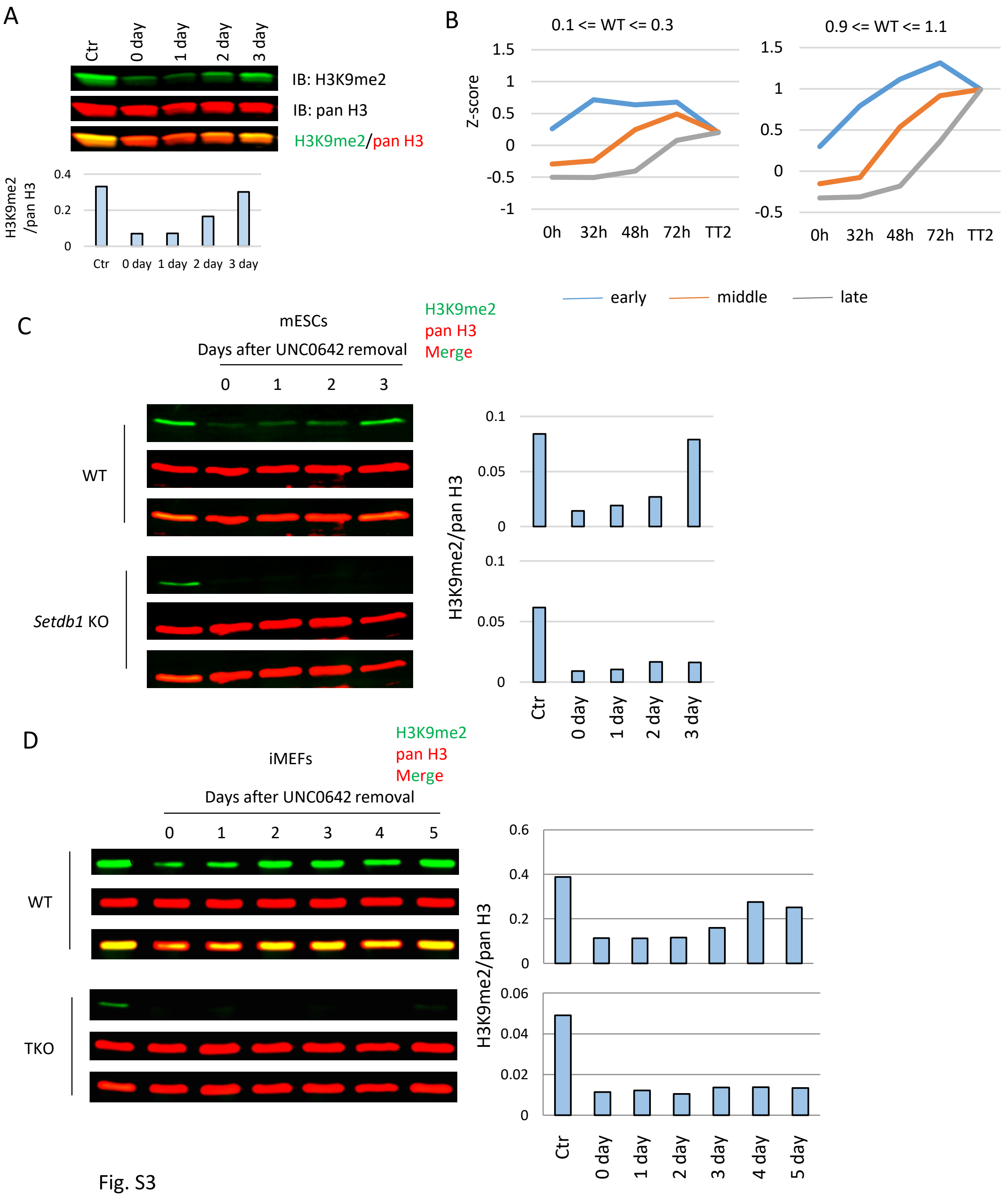

Fig. S3

A. H3K9me2 recovery after the UNC0642 treatment analyzed by western blotting. H3K9me2 is almost recovered three days after UNC0642 removal. Bottom graph shows the relative amount of H3K9me2 to pan-H3.

B. H3K9me2 recovery in each class. The left and right graphs use regions with Z-scores between 0.1 and 0.3 and between 0.9 and 1.1 in WT mESCs, respectively. In both cases, the regions that show faster H3K9me2 recovery tend to harbor a higher H3K9me2 level after the UNC0642 treatment.

C. H3K9me2 recovery after the UNC0642 treatment in *Setdb1* KO mESCs analyzed by western blotting. No clear H3K9me2 recovery is observed in *Setdb1* KO mESCs. Bar graph at bottom represents relative H3K9me2 to pan H3. Bar graph on the right represents relative H3K9me2 to pan H3.

D. H3K9me2 recovery after UNC0642 treatment in TKO iMEFs analyzed by western blotting. No clear H3K9me2 recovery is observed in TKO iMEFs. Bar graph on the right represents relative H3K9me2 to pan H3.

Fukuda et al. Fig. S4

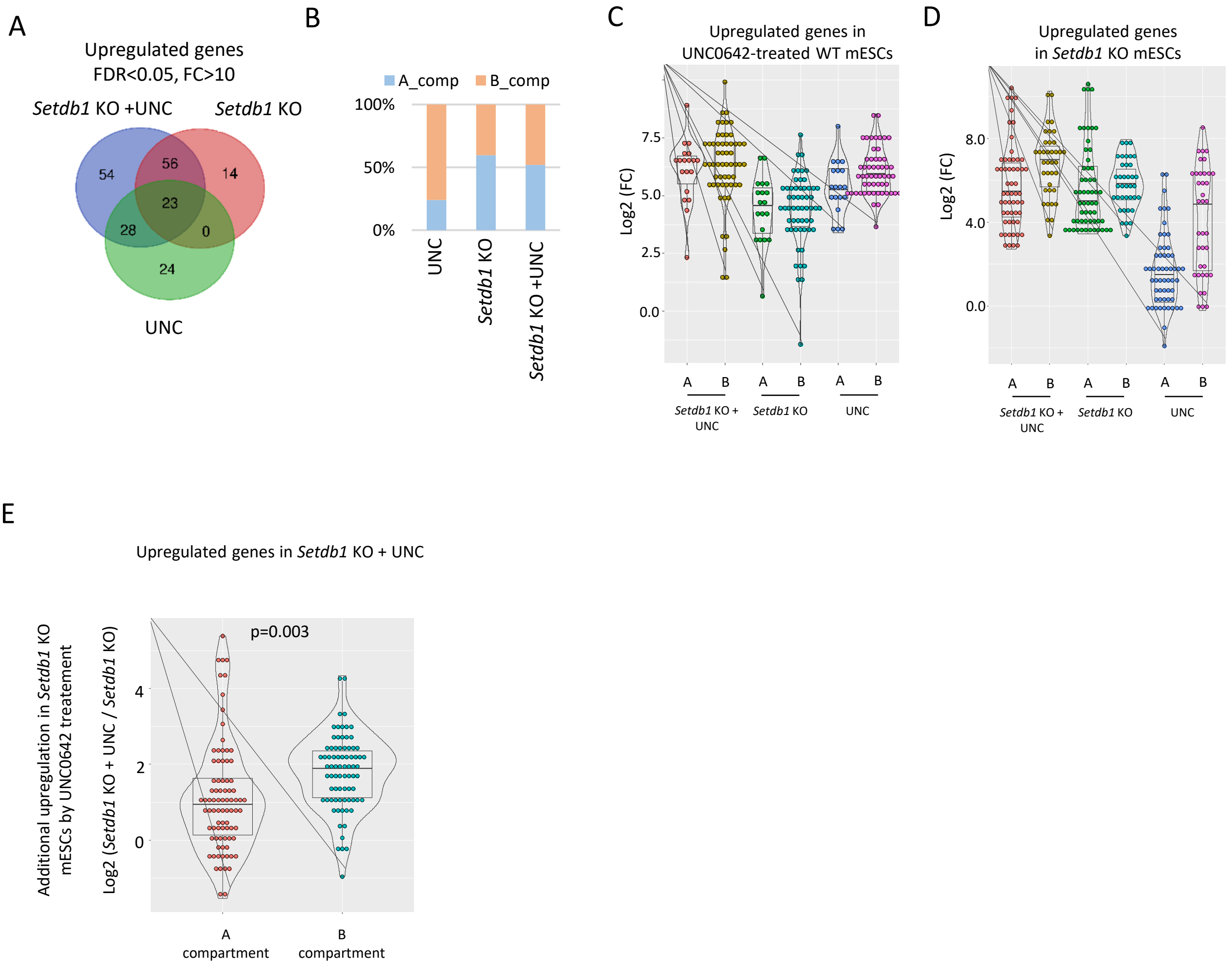

Fig. S4

A. Venn diagram of upregulated genes in each condition.

B. Distribution of upregulated genes in A/B compartment. Most of upregulated genes in UNC0642-treated mESCs are in the B compartments.

C. Violin plots of log2 fold changes in upregulated genes in UNC0642-treated mESCs. Genes in the A and B compartments are separately analyzed. Most of upregulated genes in UNC0642-treated mESCs both in the A and B compartments are derepressed also in *Setdb1* KO mESCs

D. Violin plot of log2 fold change of upregulated genes in *Setdb1* KO mESCs. Genes in A and B compartments are separately analyzed. Upregulated genes in *Setdb1* KO mESCs in the B compartments are more upregulated in UNC0642-treated mESCs than those genes in the A compartments.

E. Violin plot of log2 fold change of upregulated genes in UNC0642-treated *Setdb1* KO mESCs between *Setdb1* KO mESCs and UNC0642-treated *Setdb1* KO mESCs. Genes in A and B compartments are separately analyzed. Those genes in the B compartments are more upregulated by the UNC0642 treatment in *Setdb1* KO mESCs than those in the A compartments. P-value was calculated by student T-test.

.

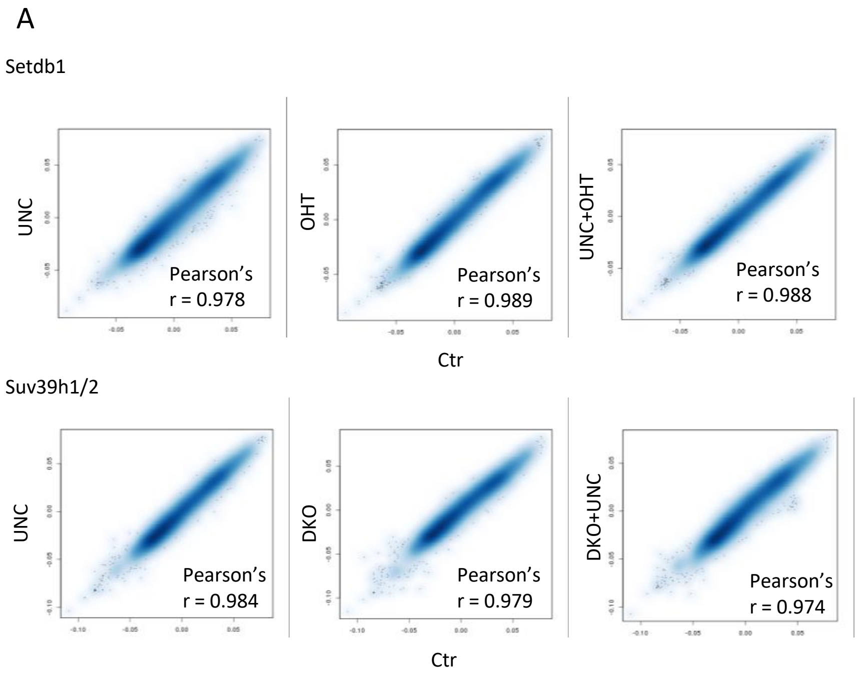

Fig. S5

A. Scatter plots of compartment score comparison between control cells and each condition.
